# Vimentin intermediate filaments undergo irreversible conformational changes during cyclic loading

**DOI:** 10.1101/673673

**Authors:** Johanna Forsting, Julia Kraxner, Hannes Witt, Andreas Janshoff, Sarah Köster

## Abstract

Intermediate filaments (IFs) are part of the cytoskeleton of eukaryotic cells and are thus largely responsible for the cell’s mechanical properties. IFs are characterized by a pronounced extensibility and remarkable resilience that enable them to support cells in extreme situations. Previous experiments showed that under strain, *α*-helices in vimentin IFs might unfold to *β*-sheets. Upon repeated stretching, the filaments soften, however, the remaining plastic strain is negligible. Here we observe that vimentin IFs do not recover their original stiffness on reasonable time scales, and we explain these seemingly contradicting results by introducing a third, less well-defined conformational state. Reversibility on the nanoscale can be fully rescued by introducing crosslinkers that prevent transition to the *β*-sheet. Our results classify IFs as a nano-material with intriguing mechanical properties, which is likely to play a major role for the cell’s local adaption to external stimuli.

---

The mechanical properties of biological cells are defined by the cytoskeleton, a composite network of microtubules, actin filaments and intermediate filaments (IF).^1,2^ Although the exact division of labour among the three filament types is still not fully resolved,^1,2^ there is ample evidence that IFs are the load bearing elements when cells are subjected to external tensile^3,4^ or compressive^5^ stress. During embryogenesis and tissue formation, in particular, cells undergo dramatic changes in shape and size. The force scales expected for cellular processes lie between single motor protein forces of a few pN, which can be measured by FRET sensors,^6^ and the collective forces of several nN measured for whole cells, as determined, e.g., by traction force microscopy.^7^ In order to withstand strong transitions, cells show reversible superelasticity, which is linked to their IF network.^3^ In order to achieve the required material properties for IFs, nature applies design principles on the nanoscale distinct from human engineering solutions and instead relies on self-organization and structural hierarchy. As a consequence, IFs stand out among the cytoskeletal filaments by their high flexibility^8,9^ and enormous extensibility.^10–13^

Within the IF family, vimentin is typical for cells of mesenchymal origin.^14^ Like all cytoskeletal IFs, vimentin monomers comprise an *α*-helical rod domain with intrinsically unstructured head and tail domains.^15^ The monomers assemble following a hierarchical pathway resulting in filaments with laterally and longitudinally arranged monomers (Fig. 1a).^16,17^ Theoretical considerations,^12,13^ molecular dynamics simulations^18,19^ and X-ray diffraction studies^20^ have shown that the intriguing tensile properties of IFs originate from conformational changes on different levels of the hierarchical filament structure.

**Figure 1:**
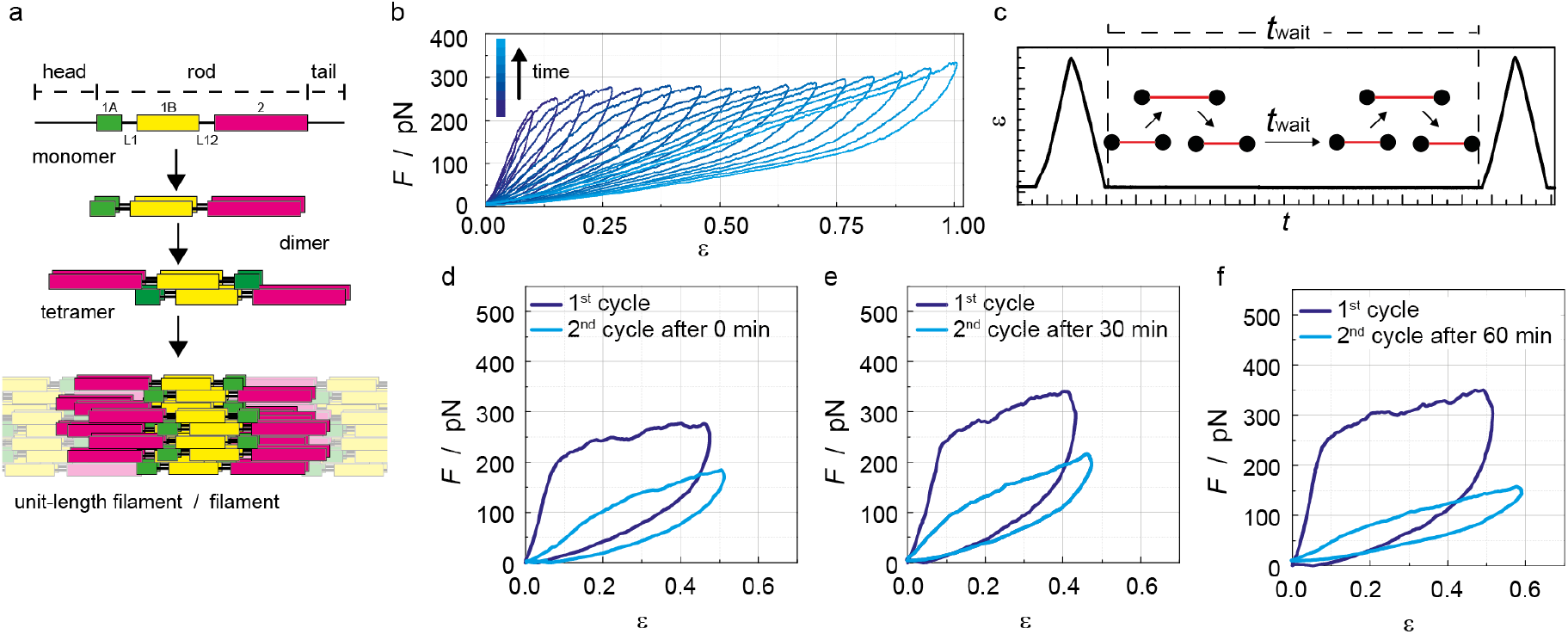
Cycle experiments on untreated vimentin IFs. **a** Schematic overview of the vimentin assembly pathway from the monomer to the mature filament, via the parallel coiled-coil dimer, the half-staggered anti-parallel tetramer and the unit-length filament that on average consists of eight tetramers. **b** Vimentin filament stretched to increasing distances with each cycle (for a filament stretched for several times to the same distance see Supplementary Fig. S1), **c** Sketch of the experimental protocol for stretching cycles including waiting time *t*_wait_. **d**, **e**, **f** Examples for force-strain data from experiments with different waiting times *t*_wait_. **d** *t*_wait_ = 0, **e** *t*_wait_ = 30 min and **f** *t*_wait_ = 60 min. For examples of all values of *t*_wait_ and a box plot diagram of the input energy ratios see Supplementary Fig. S2.

Upon stretching, the *α*-helical parts of the vimentin monomers and, as a consequence, also the dimeric coiled coils unfold, leading to an elongation of the filament. There is evidence from studies on vimentin and other IFs that the unfolded strands at least partly form *β*-sheets.^18–20^ As the measured forces during relaxation of the filament are greater than zero and we previously observed only little plastic deformation upon stretching and relaxing,^13^ it was concluded that this conformational change is reversible and that relaxed vimentin IFs eventually return to the *α*-helical state.^13^ In the same study it was observed that, when repeatedly stretched, vimentin IFs become softer with every cycle, indicated by a decrease of the slope in the force-strain curve (Fig. 1b).^13^ Monte Carlo simulations suggest that this behavior is due to the presence of mixed conformational states of parallel monomers:^13^ Fewer and fewer *α*-helical monomers survive, rendering the filament softer. These observations raise the question if and on which time scales a complete recovery of the initial mechanical properties, connected to refolding into *α*-helices, occurs.

Here, we find that, surprisingly, the initial tensile behavior cannot be recovered on reasonable time scales suggesting that this reversibility on the nanoscale is not achieved, although the filaments show negligible excess strain after relaxation. However, crosslinking of the filaments leads to a restoration of the initial tensile behavior. We present an attempt to disentangle the seemingly contradicting results towards a comprehensive understanding of the intricate mechanical behavior of IFs. We propose a consistent mechanistic model for the observed phenomena that requires to include a third, disordered, conformational state, and postulate that the native system cannot return to the *α*-helix, which becomes less populated with each loading cycle.

In order to examine the mechanical recovery after stretching of vimentin filaments we perform a straight-forward optical trapping experiment: Between two stretching cycles the filament is kept at its initial contour length for a varying time *t*_wait_ (Fig. 1c). By this procedure, conformational changes caused by straining the filament may be reversed and the filament is allowed to return to its equilibrium conformation. The first stretching cycle shows the characteristic non-linear force response described previously^13^ (Fig. 1d-f, dark blue), comprising an initial linear force increase, followed by a force plateau and eventually a pronounced hysteresis upon relaxation to zero strain. Note that in all experiments shown here, we pull the filament into the plateau region but not beyond. When the filament is stretched again (Fig. 1d-f, light blue), we observe softening, but almost reach zero strain at zero force. Furthermore, the hysteresis decreases, supporting that full recovery of the initial conformation is not achieved. However, the relaxation curve of the first cycle always lies below the stretching curve of the second cycle, indicating partial reversibility. The ratio of the input energies required to stretch the filament the first and second time shows that the degree of reversibility is independent of *t*_wait_ (Supplementary Fig. S2) even after 1 h. From these experiments, we conclude that the conformational changes occurring during the initial extension of the filament are microscopically not reversible on reasonable experimental time scales.

An easy way to limit the degrees of freedom upon unfolding is by introducing permanent crosslinkers, *i.e.* glutaraldehyde (GA, 5 C-atoms) and paraformaldehyde (PFA, 1 Catom), and thereby increasing the refolding probability along the same path on which unfolding occurred. Both aldehydes may form covalent bonds with amine groups through imine formation, in particular targeting accessible lysine residues. The longer the crosslinker, the more links are expected to form.

Qualitatively, the force-extension curves are not altered by crosslinking and show the same characteristic force plateau as found for untreated filaments, albeit shifted to higher forces (Fig. 2a). For native, untreated vimentin IFs the origin of the force plateau was previously assigned to an increase of the contour length of the filament due to force-induced conformational changes.^12,13,19^ The conservation of the curve shape after crosslinking indicates that unfolding of *α*-helices in the vimentin monomer is still possible after crosslinking with either PFA and GA. The relaxation curves of GA-crosslinked filaments, however, show a distinctly different progression compared to the untreated filaments with a force plateau and an inflection point, similar to the extension curve, albeit at a lower overall force. Force-clamp data of untreated and GA-crosslinked filaments (Fig. 2b, c) confirm that a much higher force is needed to extend the crosslinked filaments. Whereas the untreated filaments can be stretched up to strains higher than 1.0 at 250 pN, GA-crosslinked filaments stay at strains below 0.1 even after two hours at this force level. The force needed to stretch GA-crosslinked filaments to strains of about 0.8 is most likely around 400 pN.

**Figure 2:**
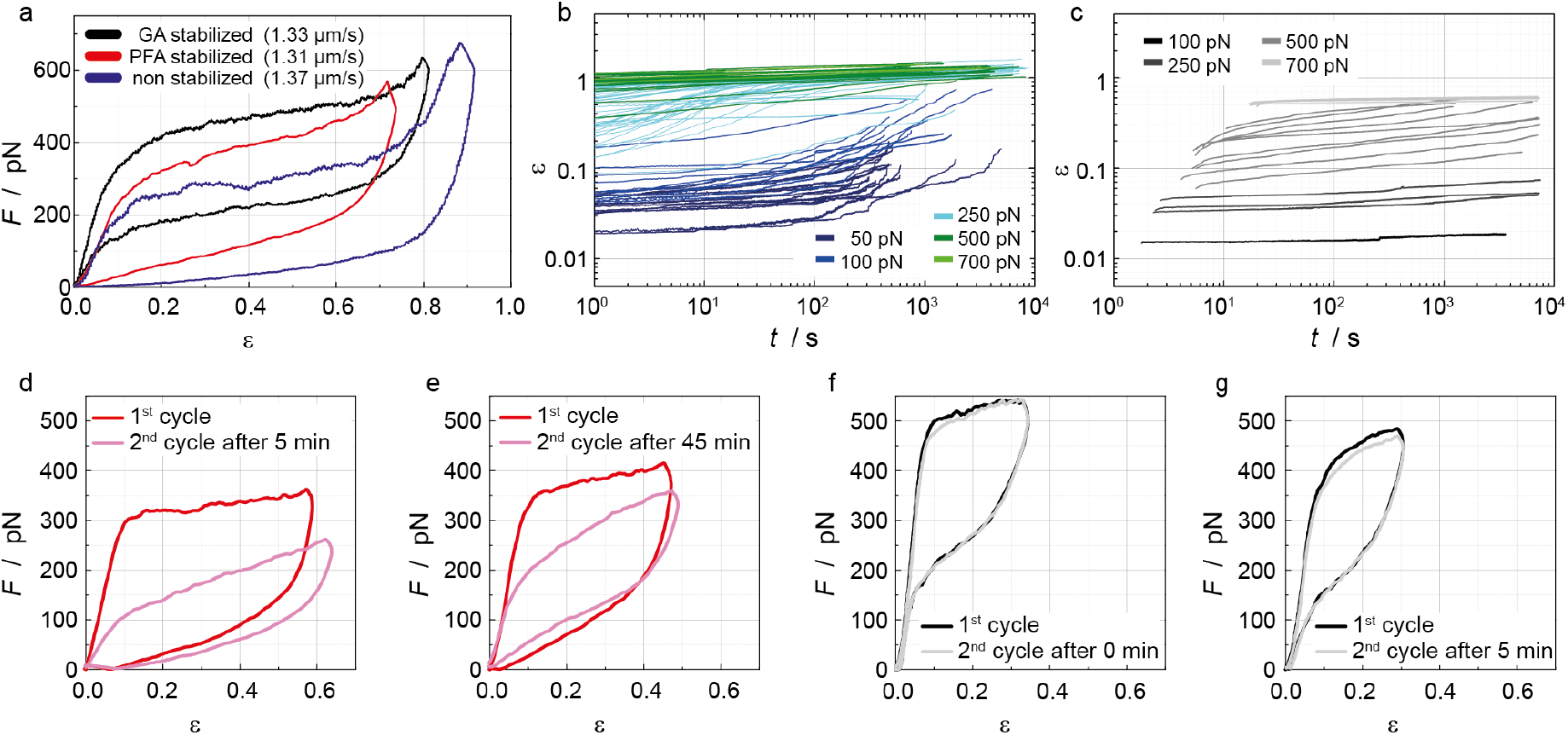
Recovery experiments with crosslinked IFs. **a** Typical force-strain cycles of untreated (blue), PFA-crosslinked (red) or GA-crosslinked (black) filaments. **b** Force clamps of untreated vimentin IFs ranging from 50 to 700 pN, presented as a log-log plot of strain versus time. ^13^ **c** Force clamps of GA-crosslinked vimentin IFs ranging from 100 to 700 pN, presented as a log-log plot of strain versus time. **d**, **e**, Examples for recovery experiments performed with PFA-crosslinked vimentin IFs with *t*_wait_ = 5 min (d) and *t*_wait_ = 45 min (e). For examples of all values of *t*_wait_ and a box plot diagram of the energy ratios for PFA crosslinked filaments, see Supplementary Fig. S3. **f**; **g** Examples for recovery experiments performed on GA-crosslinked vimentin IFs with *t*_wait_ = 0 min (f) and *t*_wait_ = 5 min (g). For examples of all values of *t*_wait_ and a box plot diagram of the energy ratios for GA crosslinked filaments, see Supplementary Fig. S4.

Repeated force-distance cycles of crosslinked filaments reveal striking differences to untreated IFs concerning the reversibility of unfolding. When PFA-crosslinked filaments are stretched again after *t*_wait_ = 5 min, they behave similar to untreated filaments (Fig. 2d). However, for *t*_wait_ = 45 min, there is a clear recovery towards the initial curve shape (Fig. 2e). By contrast, GA-crosslinked filaments show identical force-distance curves in the first and the second cycle even without any waiting time (Fig. 2f,g). These observations are confirmed by the input energy ratio as a function of *t*_wait_ (Fig. 3a). Notably, at low strain the GA-crosslinked filaments behave fully elastic, *i.e.* at small strains the relaxation curve reproduces the initial slope (Fig. 2f,g). Thus, crosslinking with GA remodels the energy landscape of IF stretching and crosslinked filaments return to the initial conformation.

**Figure 3:**
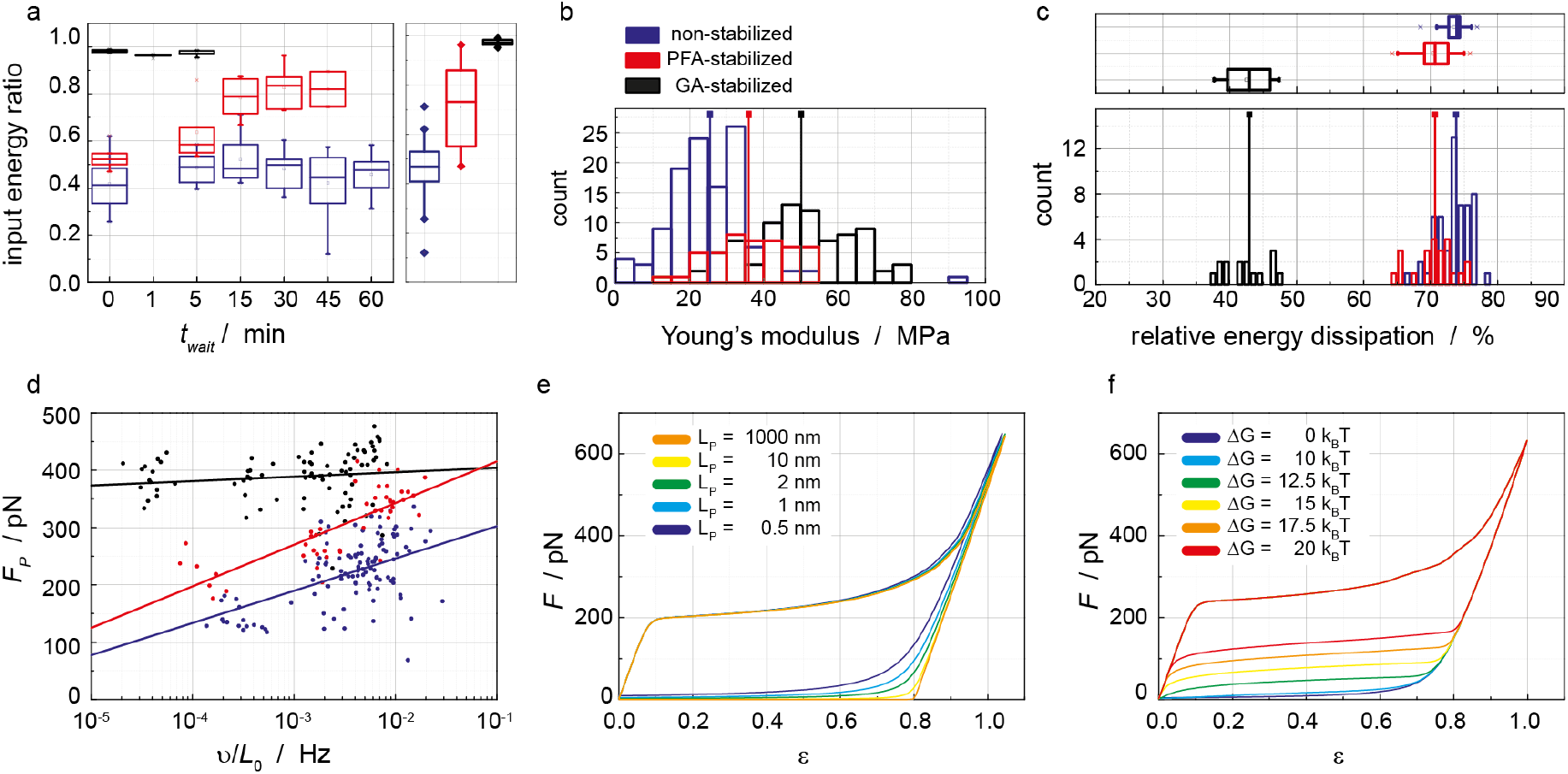
Physical analysis and numerical simulations of vimentin IF mechanics. **a** Left: ratio of input energies between the first and the second stretching event versus *t*_wait_. Right: distribution of input energy ratios for all *t*_wait_. The input energy ratio for PFA-treated filaments shows a slow increase and reaches values of about 0.85 after 30 min (red). For GA-treated filaments the input energy ratio stays constant around 1 as the first and second force-distance cycles show almost identical progressions (black). **b** Histograms of Young’s moduli of untreated, PFA- and GA-stailized IFs calculated from the initial slope (Supplementary Fig. S5) of the first force extension curve of each vimentin IF assuming a circular filament cross-section and a filament radius of 5 nm. Legend and color codes apply to a,c, and d as well. **c** Bottom: histograms of relative dissipated energies for untreated, PFA- and GA-crosslinked filaments. Top: distributions of dissipated energy. **d** Force at the onset of unfolding (*i. e.* the beginning of the plateau region) versus the normalized loading rate (Supplementary Fig. S5). **e** Simulation of force-strain curves according to Block et al. ^13^ without any back-reaction but with a decreasing persistence length for the retraction part of the curves. **f** Simulation of force-strain curves according to Block et al. ^13^ with back-reaction for different ∆G.

These results raise the question whether there exists a fundamental difference between the conformational changes during stretching for crosslinked and untreated filaments. The Young’s modulus (Fig. 3c) already shows a clear trend. It increases from 26 MPa to 36 MPa and eventually to 50 MPa for untreated (blue), PFA-crosslinked (red) and GA-crosslinked IFs (black), respectively, indicating a strong effect caused by crosslinking. Fig. 3b shows histograms of the energy dissipated during the first stretching-relaxing cycle, normalized by the input energy. For PFA-treatment (red), similar values (*∼*70-80%) to untreated filaments (blue) are found,^13^ whereas GA treatment (black) drastically decreases the energy dissipation to only *∼* 45%.

To examine the energy landscape associated with folding and unfolding, the force at the onset of the plateau is determined and plotted against the loading rate *v* normalized by the initial filament length *L*_0_ (Fig. 3d). Whereas the plateau forces for PFA-crosslinked and untreated filaments show a similar velocity dependence, albeit with slightly higher plateau forces for the crosslinked filaments, the GA-crosslinked filaments show very high plateau forces around 400 pN that are almost independent of the loading rate. We have shown previously that the force-extension curve of vimentin can be modelled with an elastically coupled two-state-model.^12^ This model predicts a logarithmic scaling of the plateau force with pulling velocity (see Supporting Information, equation 5). For the untreated filaments we find a potential width *x*_*u*_ = 0.17 *±* 0.02 nm (mean *±* st.d.) and a zero-force reaction rate of *k*_0_ = (5 *±* 4) ⋅c> 10^*−*7^ s^*−*1^ in good agreement with the parameters found previously by directly fitting to individual force-strain curves.^12^ Treatment with PFA leads to an only slightly decreased potential width of *x*_*u*_ = 0.13 *±* 0.02 nm and zero-force reaction rate of *k*_0_ = (2.5 *±* 2) ⋅c> 10^*−*7^ s^*−*1^. By contrast, GA drastically alters the energy landscape. The reaction is slowed down dramatically, *k*_0_ = (1 *±* 8) ⋅c> 10^*−*54^ s^*−*1^, which is accompanied by an enormous increase of the potential width, *x*_*u*_ = 1.2 *±* 1 nm. This huge change probably indicates that the molecular elongation mechanism is fundamentally different after GA-treatment. Therefore, it is remarkable that basic characteristics of the tensile behavior like the high extensibility and the force plateau are largely conserved after crosslinking.

We can only speculate about the molecular origin of the observed changes due to crosslinking. However, a previous crosslinking study employing disulfosuccinimidyl tartrate (DST), which is only one carbon atom shorter then GA, led to a dense network of crosslinks in vimentin IFs.^21,22^ It has been shown previously that the formation of intramolecular cross-links is highly unlikely. Downing^22^ showed that crosslinking between two lysine groups that are both within an *α*-helical region is unusual. This is true within dimers as well as between dimers as steric hindrance prevents linkage between lysines in identical sequence locations in the *α*-helical regions of parallel chains. As a consequence, intermolecular crosslinks are predominately found in random coil conformations and beta sheets. It is expected that these crosslinks prevent the necessary rotation of the peptide chain for the *α*-helices to completely unfold, so we regard them as pinning points for the protein conformation, which ensure the presence of a reasonably close starting point for the reformation of the *α*-helices.

Molecular dynamics simulations have shown that coiled coils need a critical length of about 40 amino acids in order to be able to undergo an *α*-*β*-transition,^23^ which is in good agreement with the dramatically increased stability we observed for the short *α*-helical segments between the crosslinks. Furthermore, we assume that, when the *α*-helices in GA-crosslinked filaments unfold, parallel helices have to unfold simultaneously since they are covalently connected. This is in contrast to the untreated filament, where computational models have predicted that parallel unfolding is initiated by stochastic opening of a few monomers which leads to loss of stability of all remaining parallel helices,^13^ similar to what has been observed in binding clusters.^24,25^ Using the analogy to binding clusters, this can be regarded as an extreme case of shared loading which might explain the huge increase of the potential width.^26,27^

The relaxation force curve of untreated vimentin IFs as shown in Fig. 2a, blue, decays almost linearly from a restoring force around 100 pN at a strain of 0.8 to zero force at strains below 0.1. This positive restoring force was earlier interpreted as a sign of reversibility of the underlying molecular process. This can be illustrated by modelling the force extension curve using a two-state-model for the filament elongation introduced earlier.^13^ If we switch off the back-reaction in the model (Fig. 3e) the force drops to zero already at large strains around 0.8 if we assume that the persistence length of the filament stays constant at 1 *µ*m upon stretching (orange). If, however, the persistence length is decreased dramatically upon stretching, a force progression closer to the experimental relaxation curve is recovered (green, blue, purple). This is plausible, since a thinning of IFs upon stretching has been observed^10,11^ and would lead to smaller persistence lengths. Furthermore, since coiled coils are particularly rigid, it is likely that the filament gets more flexible when being unfolded. Indeed, a similar mechanism with a drastic decrease in persistence length upon unfolding was proposed by Minin et al. for the *α*-*β*-transitions in simpler coiled-coil peptides.^28^ In contrast to an earlier interpretation, we conclude that entropic elasticity of the protein chain is sufficient to explain the restoring force upon filament relaxation for untreated vimentin filaments.

The force plateau and inflection point in the relaxation curves of GA-crosslinked filaments (Fig. 2a, black), are reproduced by the two-state-model when increasing the energy difference ∆*G* between the shorter and the longer state and thereby the reversibility of the transition (Fig. 3f). This indicates that the positive restoring force of GA-crosslinked filaments originates from the refolding enthalpy of the *α*-helices. Interestingly, a similar increase in reversibility might also be achieved by a symmetrization of the energy landscape.

The strongest argument for the proposed reversibility of vimentin elongation was the observation that repeated cycles always start at nearly the same strain (Fig. 1a) and that only very little residual plastic strain of vimentin was observed (Supplementary Fig. S6). The softening of filaments without residual strain was explained by mixed states – *α*helical and elongated – in the parallel monomers. However, the observed force plateau requires all parallel monomers in a unit length filament to elongate in a cascading manner.^13^ Thus the softening with repeated cycles can only be explained when there is a way for elongated monomers to return to a shorter effective length, which seems to contradict the non-reversibility we observe here.

To resolve the contradiction, we propose a model for which the transition between the shorter and the longer state is not a one-step process as assumed in the two-state-model proposed previously,^12^ but consists of three distinct states (Fig. 4a). When a force is applied to the filament, the *α*-helical coiled coils unfold (black arrow) and the polypeptide chains assume a random coil. This unfolding can be described as a transition along a reaction coordinate orthogonal to the applied force. Due to its high energy cost this reaction does not occur spontaneously but only at high loads around 200 pN, *i.e.* the plateau force. The unfolded chain can elongate in response to the applied strain leading to a reversible extension of the monomer (blue arrow). This elongation has presumably not only an entropic cost to extend the semiflexible polypeptide chain, but also an enthalpic component due to the breaking and formation of hydrogen bonds, rendering the energy landscape of the unfolded state rather rough. This roughness of the energy landscape might indeed give rise to the viscoelastic relaxation behavior investigated earlier, where we find a power law behavior.^13^ When the random coil is elongated, hydrogen bonding between parallel polypeptide chains can lead to the formation of *β*-sheets (red arrow). This process is presumably rather slow, since the polypeptide chains need to assume the exact right geometry, and irreversible, due to the high energy difference. When the filament is subsequently relaxed, only the peptide chains that were not able to form a *β*-sheet will return to a shorter contour length. Since the relaxed unfolded polypeptide chain can assume many possible conformations, reformation of *α*-helices is highly unlikely (dashed arrow).

**Figure 4:**
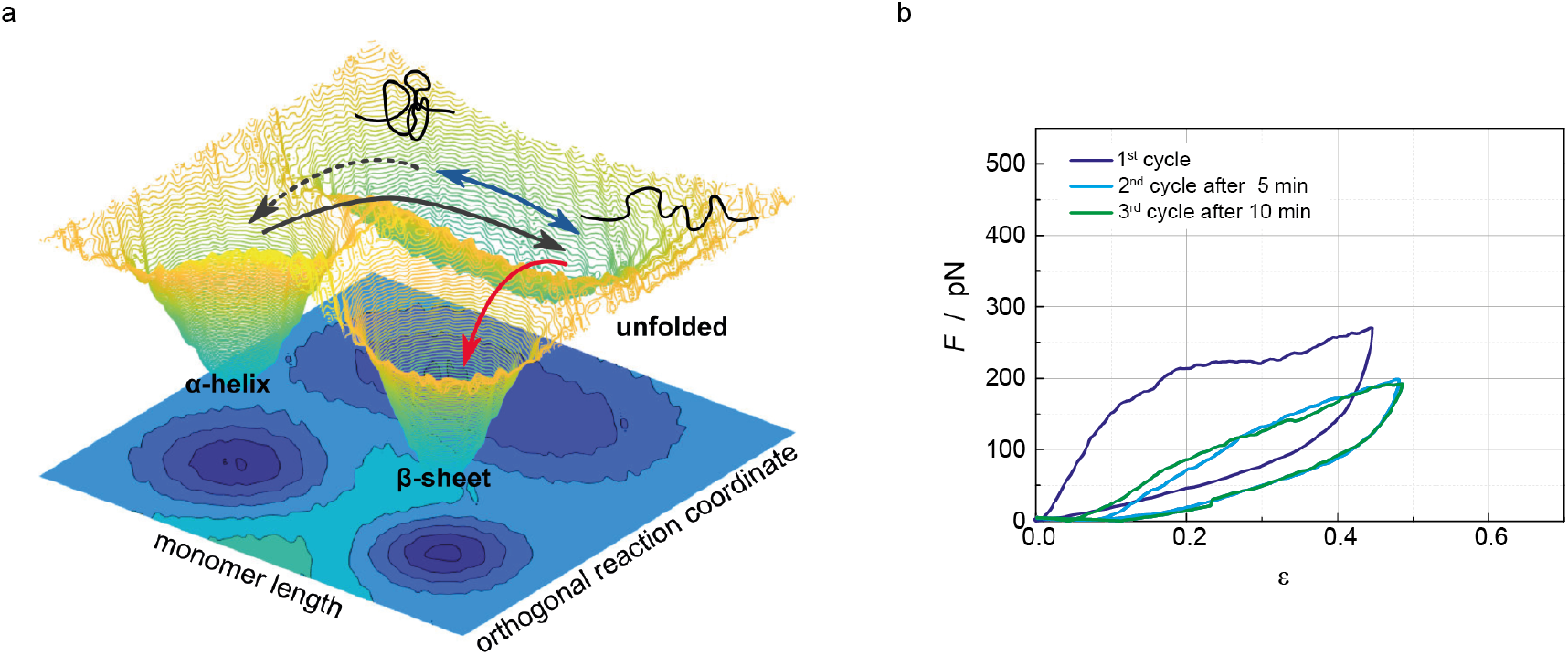
Introduction of a third conformational state during vimentin IF stretching. **a** Energy landscape illustrating the proposed model. Besides the two well-defined *α*-helical and *β*-sheet conformations, there exists a less defined unfolded conformation along an orthogonal reaction coordinate. When a force is applied, the *α*-helix can unfold (black arrow) and the unfolded random coil can be stretched (blue arrows). **b** An untreated vimentin IF is cycled three times with *t*_wait_=5 min between the cycles. Whereas the force-strain behavior of the second cycle differs considerably from the first one, the data for the third cycle are almost identical to the second one.

Our model is supported by an experiment, where we cycle an untreated filament three times with a waiting time of *t*_wait_= 5 min between the cycles. Whereas the data for the second cycle (light blue) differ from the first one (dark blue), in line with the data shown in Fig. 1, the third cycle (green) progresses in an identical manner to the second one. In our model energy landscape (Fig. 4a) we interpret this as follows: upon first stretching, the *α*-helices are unfolded into random-coils, which are subsequently elongated. Relaxation and repeated stretching leads to “shuffling” between the short and the long random-coil state and thus to identical force-strain curves.

This model is sufficient to justify why the filament always returns to essentially the same contour length, while also explaining the observed irreversibility. Apparently, PFA crosslinked polypeptide chains are kept close enough to the starting conformation that slow refolding of the *α*-helices is observed, leading to a slow recovery of the initial force extension curve. By contrast, the energy landscape of the GA-crosslinked filaments is strongly altered, and rather resembles the idealized two-state-model, with an energetically heavily favored shorter state and a spontaneous back-reaction from the elongated state. From our measured force-strain cycles (Fig. 2a, black) and the proposed model, one may draw an analogy to pseudoelastic (also referred to as superelastic) shape-memory alloys, which, when heated after being strained, return to a previous shape.^29^ Of note, however, in our system, we do not observe a phase transition, as required for pseudoelasticity.

By employing controlled *in vitro* experiments, we are able to precisely tune mechanical strain-response of vimentin IFs to applied stress. Untreated IFs can be stretched to high strains and largely recover their original length upon relaxation, *i.e.* only neglectable residual strain is observed. As the filaments soften with each subsequent cycle and the original stiffness is not recovered on reasonable experimental time scales, we examine the energy landscape of strained IFs in detail. Our experiments lead to the conclusion that besides the *α*-helix and the *β*-sheet a third – presumably random coil – conformational state must exist, and that subunits are not able to return to their initial state after stretching. This irreversible transformation is avoided when introducing permanent crosslinkers into the system and thereby reducing the degrees of freedom upon stretching.

The irreversibility of the conformational changes we observed here might indicate that vimentin filaments that were strained in the past are not as capable to protect the cellular integrity. This implies that cells need a mechanism to “repair” strained vimentin.

This reversibility may be achieved by either subunit exchange along the filaments,^30–32^ which is, however, very slow, or by disassembly and reassembly of filament structures as observed *e.g.* for keratin networks.^33,34^

Interestingly, the microscopic reversibility and the increased stiffness we observed as a result of GA-fixation seem to be advantageous mechanical properties for cells that are heavily subjected to external strains. For keratin IFs in maturating keratinocytes, controlled covalent crosslinking of subunits by disulfid bond formation is indeed observed and might be a possible avenue for cells to tune their resistance to external forces.^35,36^ Interestingly, vimentin was identified as a substrate for proteins introducing covalent crosslinks between different monomers, specifically transglutaminases, in two human tissues where structural integrity is paramount, arteries and lens.^37,38^ Like the crosslinkers used in our work, transglutaminases target primary amines suggesting they might alter the energy landscape in a way similar to what we show in our *in vitro* system.

## Methods

### Vimentin production and purification

The production and purification of recombinant human vimentin C328A with additional amino acids GGC at the C-terminus for force clamp and multiple-cycle-measurements was performed as described previously.^13^ For all other experiments the protein purification was performed similar to the protocol described previously^13^ but with several changes to the buffers used, as described in the Supplementary Information.

### Vimentin and bead functionalization

Vimentin was labeled, reconstituted, and assembled according to previously published protocols. Labeling with ATTO647N (ATTO-Tech GmbH) and biotin-maleimide (Jena BioSciences GmbH) was performed according to Block et al.^12^ and Winheim et al.^39^ Reconstitution and assembly were performed as described by Block et al.^13^

Carboxylated polystyrene beads were functionalized according to Janissen et al.^40^ as described in Block et al.^13^

### Optical tweezers measurements

Optical tweezers measurements were performed using a commercial instrument (C-trap, Lumicks, Amsterdam, Netherlands), combining optical tweezers with a microfluidic setup and confocal microscopy (Supplementary Fig. S7), similar to what was described previously.^13^ In short, a fresh pair of beads was captured for each experiment in the bead channel. A vimentin IF was captured in the vimentin channel and bound to the second bead in the buffer (100 mM KCl in 2 mM phosphate buffer, pH 7.5) channel. Parts of the measurements were performed with filaments bound to the beads by maleimide-chemistry and parts of the experiments with filaments bound via biotin-streptavidin. The covalent attachment of the filaments via maleimide-groups enables us to introduce waiting times between the filament stretching cycles. For crosslinking experiments the captured vimentin IF was moved to channel 4 (Supplementary Fig. S7) filled with buffer containing either PFA (0.12 % (v/v)) or GA (0.5 % (v/v)). Before a force-distance curve was recorded, the filaments were always moved back to the channel that only contains buffer (channel 3, Supplementary Fig. S7). Additionally, a prestrain of 5 pN was imposed for all recovery experiments. This prestrain prevents the filaments from wrapping around the beads during the waiting time as well as the loss of the filament during that time. Before the actual measurement, this value was set to 0 pN and the thus defined prestrained length of the filament was used as initial length *L*_0_ to calculate the dimensionless strain during stretching.

Following the measurement protocol sketched in Fig. 1b, we stretched the filament beyond the linear regime into the characteristic plateau region, where unfolding of the *α*-helices occurs. Subsequently, we relaxed the filament and, after a defined time *t*_wait_ *≤* 1h, stretched the filament again to the same trap position as in cycle one. The length of the filament and the applied force were recorded over the full experiment.

### Data Sets

For the untreated filaments 57 individual recovery experiments were performed. The total number of recovery measurements for GA-crosslinked filaments was 15 and for PFA-crosslinked filaments it was 23. For the force-clamp experiments of untreated filaments about 100 measurements were performed. The number of force-clamps of GA-crosslinked filaments was 18.

### Data analysis

All data were converted to ASCII format with the measuring software (TWOM, Lumicks). The bead diameter was subtracted from the distance and the time and force data were used as measured. To calculate the energy for the first cycle, the area beneath the stretching curve up to the maximum strain was calculated by integration. As we can only control the position of the trap and not the position of the bead, the maximum strain of the second stretching is slightly higher than the maximum strain for the first stretching cycle, due to the softening of the filament. To compare the input energy that was needed to strain the filament to the same length again, the energy needed for the second stretching was calculated by integrating the area beneath the second stretching curve but only up to the maximum strain of the first stretching curve. The two energy values were then divided to obtain the input energy ratio. In order to determine the Young’s modulus, the initial slope of the force extension curve was determined by a linear fit. Only data points with forces larger than 10 pN and smaller than 80% of the plateau force and at positive strains where considered. The force at the onset of the plateau was determined by connecting the first and the last data point of an extension curve with a straight line and subtracting this straight line from the force curve. This effectively tilts the force curve, such that the beginning of the plateau can be determined as the maximum of the tilted curve. A logarithmic scaling law (see Supporting Information, equation 5) was fitted to the plateau force as a function of the pulling velocity *v* normalized by the initial filament length *L*_0_. To estimate the uncertainty of the parameters, a bootstrapping algorithm^41^ with 10000 iterations was used.

## Supporting information

Supporting Informations

## Data availability

The data that support the findings of this study are available from the corresponding authors, at, or, upon reasonable request.

## Acknowledgements

The authors thank Andrea Candelli and Jordi Cabanas Danes for fruitful discussions and technical support. The work was financially supported by the European Research Council (ERC) under the European Union‘s Horizon 2020 research and innovation program (Consolidator grant agreement no. 724932 and Laserlab-Europe grant agreement no. 654148). Further financial support was received from the Deutsche Forschungsgemeinschaft (DFG) in the framework of SFB 755 (project B7) and SFB 937 (project A17).

## Author contribution

SK conceived the project. JF and JK each performed parts of the experiments and the data analysis. HW performed parts of the data analysis, developed and implemented the models and performed all simulations. AJ and SK supervised the project. All authors wrote the manuscript.

## Supporting Information

Additional figures, detailed methods, an analytical solution for the two-state-mode and additional references^41,42^ are supplied as Supporting Information.

