## Supporting Informations for "Vimentin intermediate filaments undergo irreversible conformational changes during cyclic loading"

### Supporting Information for: ”Vimentin Intermediate Filaments Undergo Irreversible Conformational Changes during Cyclic Loading”

Johanna Forsting[1#], Julia Kraxner[1#], Hannes Witt[2#], Andreas Janshoff[3*], Sarah Köster[1*]

*
### equal contribution

[1]Institute for X-Ray Physics, University of Goettingen, 37077 Göttingen, Germany
[2]Max Planck Institute for Dynamics and Self-Organization, 37077 Göttingen, Germany
[3]Institute of Physical Chemistry, University of Goettingen, 37077 Göttingen, Germany

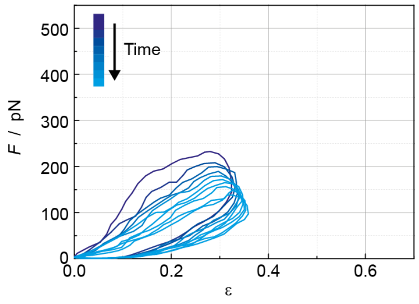


**Figure S1:** Stretching cycle experiment. A vimentin filament is stretched several times to an almost constant distance. For technical reasons, a fully constant distance is not possible.

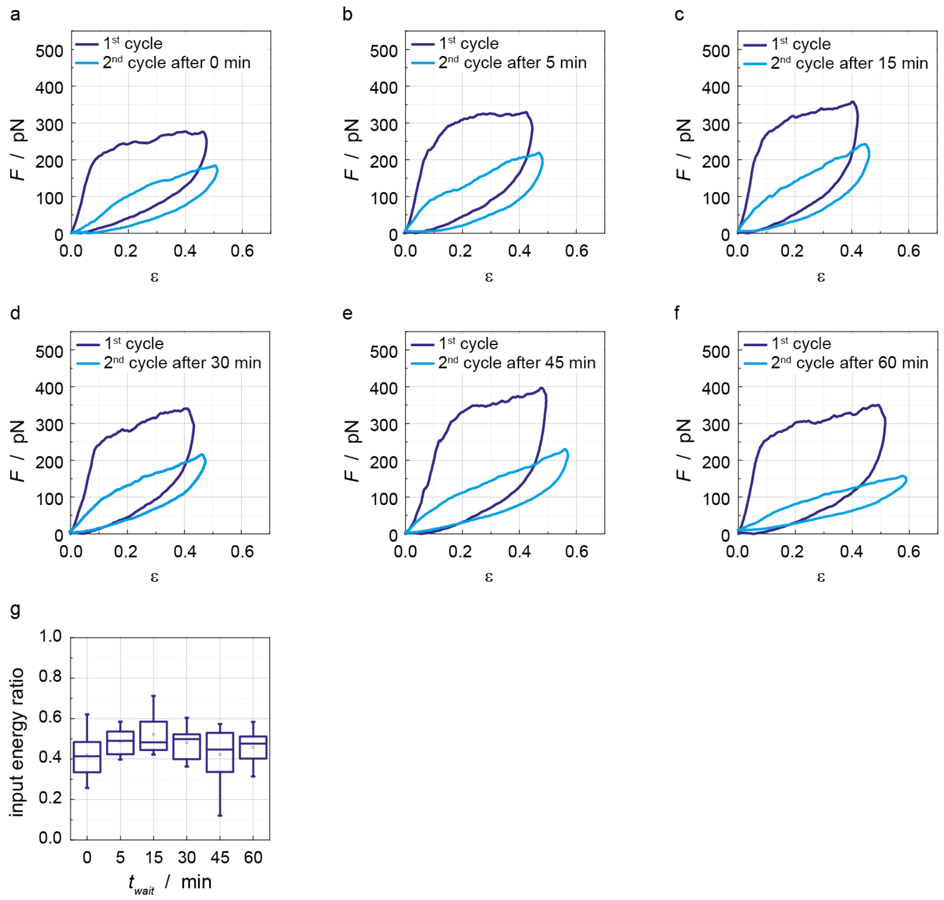


**Figure S2:** Recovery experiments with untreated vimentin filaments. Examples for force-strain data from experiments with different waiting times $t_{\mathrm{wait}}$. **a** $t_{\mathrm{wait}} = 0$ min. **b** $t_{\mathrm{wait}} = 5$ min. **c** $t_{\mathrm{wait}} = 15$ min. **d** $t_{\mathrm{wait}} = 30$ min. **e** $t_{\mathrm{wait}} = 45$ min. **f** $t_{\mathrm{wait}} = 60$ min. **g** Box plot of input energy ratios versus $t_{\mathrm{wait}}$.

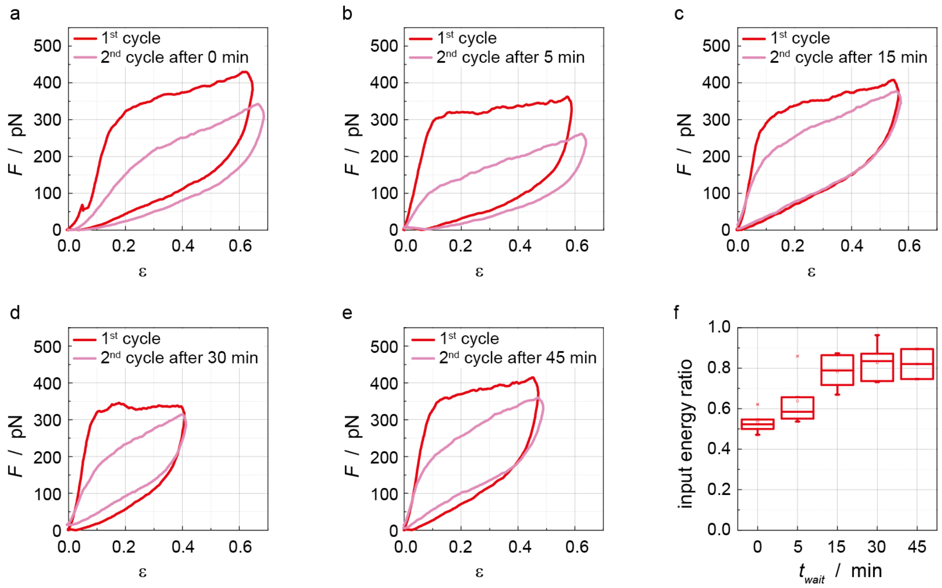


**Figure S3:** Recovery experiments with PFA-crosslinked vimentin filaments. Examples for force-strain data from experiments with different waiting times $t_{\mathrm{wait}}$. **a** $t_{\mathrm{wait}} = 0\,\mathrm{min}$. **b** $t_{\mathrm{wait}} = 5\,\mathrm{min}$. **c** $t_{\mathrm{wait}} = 15\,\mathrm{min}$. **d** $t_{\mathrm{wait}} = 30\,\mathrm{min}$. **e** $t_{\mathrm{wait}} = 45\,\mathrm{min}$. **f** Box plot of input energy ratios versus $t_{\mathrm{wait}}$.

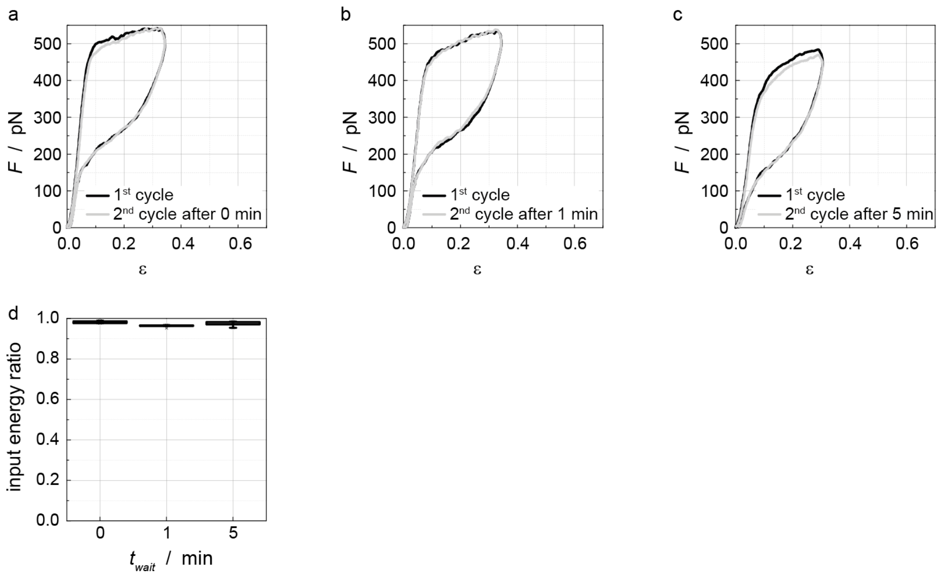


**Figure S4:** Recovery experiments with GA-crosslinked vimentin filaments. Examples for force-strain data from experiments with different waiting times $t_{\mathrm{wait}}$. **a** $t_{\mathrm{wait}} = 0\,\mathrm{min}$. **b** $t_{\mathrm{wait}} = 1\,\mathrm{min}$. **c** $t_{\mathrm{wait}} = 5\,\mathrm{min}$. **d** Box plot of input energy ratios versus $t_{\mathrm{wait}}$.

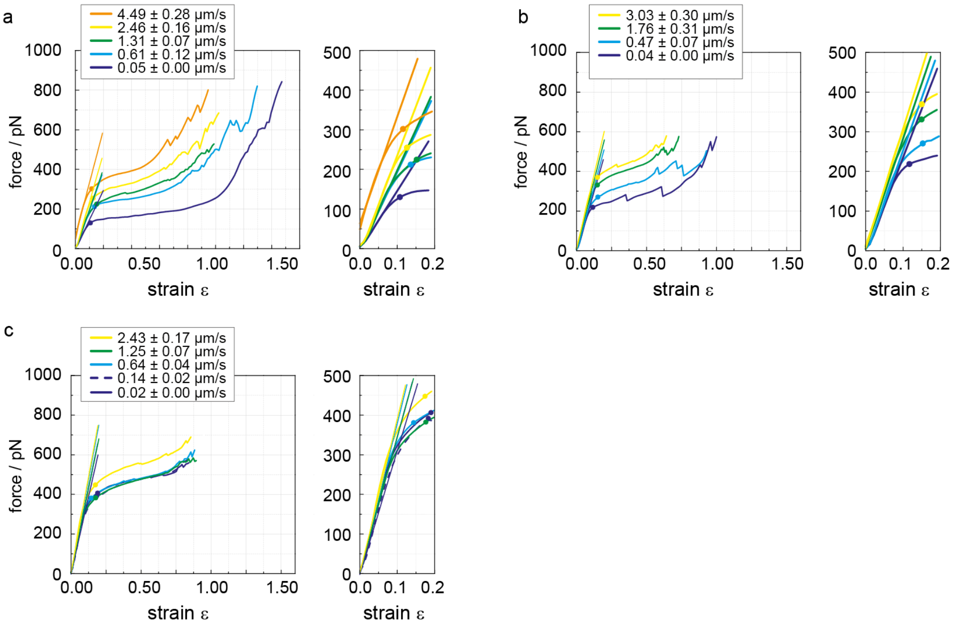


**Figure S5:** Typical force strain curves (left) and magnifications of the elastic initial regime (right) of **a** untreated, **b** PFA-crosslinked, and **c** GA-crosslinked vimentin filaments at different pulling velocities (color-coded) illustrating how the Young's modulus (linear fit) and the plateau force (dot) are quantified.

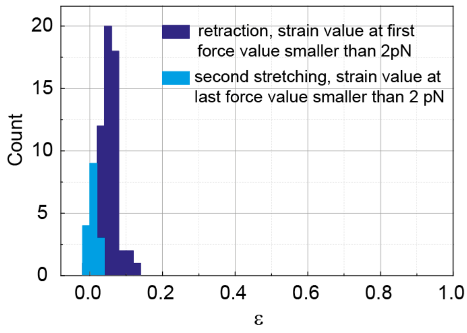


**Figure S6:** Quantification of remaining strain. When we quantify the strain an untreated filament has reached when the measured force during the retraction drops below 2 pN, we gain a distribution of strains with a median at 0.05 and a maximum at 0.12 (dark blue). The strains we observe for the last force value smaller than 2 pN for the second stretching are even smaller values (light blue). The histogram of strains for the second stretching shows a median at 0.007 and a maximum at 0.03, supporting the hypothesis that there is only neglectable plastic deformation in the filament.

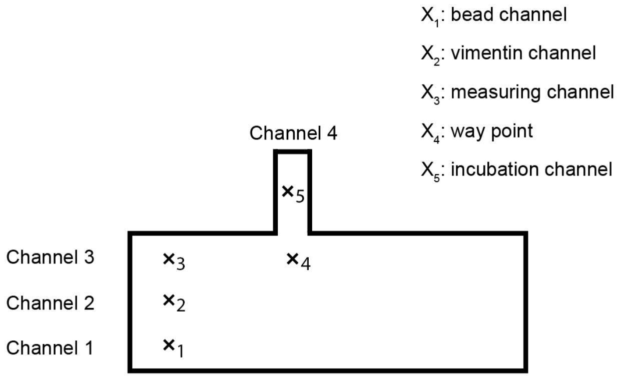


**Figure S7:** Schematic of the flow cell geometry. The marked positions $x_1$ to $x_5$ indicate where the experimental steps were performed. Two beads were captured at position $x_1$, vimentin filaments were captured at position $x_2$. Position $x_3$ was used to bind a single filament to both beads and for performing the actual measurements. Incubation with crosslinking chemicals GA and PFA was performed in channel 4, position $x_5$, while position $x_4$ was only a way point for the automated stage drive to reach channel 4 without loosing beads and filaments. Otherwise the system would move the stage the direct way from $x_3$ to $x_5$, the traps would leave the microfluidic channel and the beads and the filament would get lost at the channel wall.

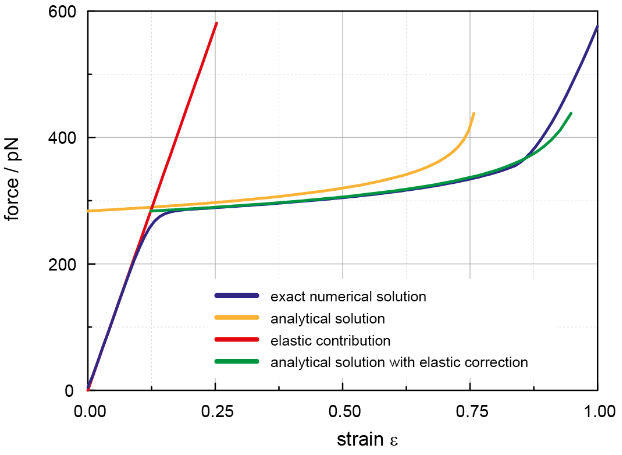


**Figure S8:** Comparison of the exact numerical solution (blue) to the analytical approximation with (green) and without (yellow) elastic correction (red). Although the stiffening at large strains is not quantitatively reproduced, the agreement in the low strain-regime that is used to predict the scaling law is good.

#### Vimentin production and purification

*Escherichia Coli* bacteria (100 $\mu$L) were thawed on ice, mixed with 2 $\mu$L plasmid solution (concentration 10 ng/$\mu$L) per 50 $\mu$L bacteria solution and cultured on an agar plate [1]. A single bacteria colony was picked and transferred into 5 mL terrific broth medium (TB, 47.6 g/L (#T0918, Sigma-Aldirch), containing glycerol (8 mL of 99.5 % glycerol (#3873, Sigma-Aldrich) per 1 L TB medium) and ampicillin (Sigma-Aldrich, added at a concentration of 1:1000) and incubated at 37° C at 220 rpm for four to six hours. Afterwards, the bacteria solution was transferred to 50 mL TB medium and allowed to grow for another four to six hours. Finally the bacteria were transferred to 1 L of TB medium and cultured over night at 37 °C at 150 rpm.

Bacteria were pelleted with a Beckmann Centrifuge (Beckmann Coulter Avanti J-26 XP, rotor JS5.3) at 5000 $g$ at 4 °C for 25 min. For purification, the pellet was transferred to a cooled Douncer and kept on ice for all subsequent steps. The pellet was dissolved in 16 mL lysis buffer (50 mM TRIS (Carl-Roth GmbH), pH 8.0, 25 % w/v saccharose, 1 mM EDTA (Carl-Roth GmbH), 1 mM Pefabloc SC® (Carl-Roth GmbH), cOmplete™ tablets (1 per 50 mL, Sigma-Aldrich)). 4 mL lysozym (Roche Diagnostics) solution (10 mg/mL in lysis buffer) were added, the mixture was homogenized and incubated on ice for 30 min. 200 $\mu$L of 1 M $MgCl_2$ (Sigma-Aldrich), 20 $\mu$L RNAseA (10 mg/mL in TRIS buffer, pH 7.5, Roche Diagnostics), 2 $\mu$L benzonase (#71206-6, Novagen) and 400 $\mu$L 10 % NP-40 (Roche Diagnostics) were added, the solution was again homogenized and incubated for 10 min. 40 mL detergent buffer (200 mM NaCl (Carl-Roth GmbH), 1 % NP40, 1 % DOC (sodium-deoxycholat, Sigma-Aldrich), 20 mM TRIS, 2 mM EDTA, 1 mM Pefabloc SC, cOmplete™ tablets (1 per 50 mL)) were added, the mixture was again homogenized, incubated for 10 min, and then centrifuged for 30 min at 10,000 $g$ and 4 °C using a Beckmann centrifuge (J26XP, rotor JLA 16.250).

The supernatant was discarded and the pellet washed by homogenization using 4 °C cold GII-buffer (10 mM TRIS, 0.1 Vol% TritonX-100 (Carl-Roth GmbH), 5 mM EDTA, pH 8.0) complemented by 40 $\mu$L 1 M DTT (1,4 dithiothreitol, Carl-Roth GmbH) and 100 $\mu$L Pefabloc SC. The solution was incubated on ice for 10 min and then centrifuged for 30 min using the same parameters as in the step before.

The supernatant was discarded and the pellet washed by homogenization using 4 °C cold GII-KCl-buffer (1.5 M KCl (Carl Roth GmbH) in GII-buffer) complemented by 40 $\mu$L 1 M DTT and 100 $\mu$L Pefabloc SC. The solution was incubated on ice for 30 min and then centrifuged for 30 min using the same parameters as in the step before.

The supernatant was discarded and the pellet was washed by homogenization using 4 °C cold TE-buffer (10 mM TRIS, 0.1mM EDTA, pH 8.0) complemented by 40 $\mu$L 1 M DTT and 100 $\mu$L Pefabloc SC. The mixture was incubated on ice for 10 min and then centrifuged for 25 min using the same parameters as in the step before.

The supernatant was discarded and the pellet solubilized in hand-warm urea buffer. For the urea buffer 12.125 mL 9.5 M urea solution was complemented by 125 $\mu$L 1 M TRIS, 125 $\mu$L 0.5 M EDTA and 125 $\mu$L 1 M DTT, pH 7.5, and the pellet solubilized in as less buffer as possible. After homogenization the solution was finally centrifuged at 20 °C and 100,000 $g$ for 60 min, using a Beckmann ultracentrifuge (Beckmann Coulter Optima L90K, rotor Ti70). The supernatant that contains the vimentin was in some cases stored at -80 °C.

The final purification by anion and cation exchange chromatography was performed as described in Block et al.[1] without any changes.

#### Analytical solution for the two-state-model

Here we use an approximate analytical solution of the two-state-model for filament stretching. The full theory and its applicability have been described elsewhere [1, 2, 3]. In brief, every unit length filament (ULF) in the filament is assumed to be able to adopt a shorter and a longer state with a force dependent transition between them. These states do not necessarily need to be identical to defined secondary structures like the $\alpha$-helix or the $\beta$-sheet, but can be interpreted as a two-dimensional projection of the higher dimensional energy landscape shown in Fig. 4a in the main text. This model gives us the time and force dependent contour length $L_C(t, F)$ of the filament that can be combined with elastic and entropic contributions to calculate the shape of the force-distance curve (for the sake of readability, we will state neither explicit nor implicit dependencies going forward). The approach for the analytical solution is mostly adapted from Burte and Halsey [2]. The key step for the analytical solution is to ignore elastic and entropic effects such that the filament length equals the contour length $L = L_C$. When we neglect the back-reaction to the shorter conformation, which is a good approximation for the extension curve, and do not account for the independent reaction of the separate helices in the vimentin monomer, the rate equation reads

$$\frac{\mathrm{d}N}{\mathrm{d}t} = (N_0 - N)k_0 \exp\left(Fx_u/k_BT\right), \tag{1}$$

with the number of ULFs in the longer state $N$, time $t$, the total number of ULFs in the filament $N_0$, the zero force reaction rate between the shorter and the longer state $k_0$, the force $F$, the potential width $x_u$, Boltzmann's constant $k_B$ and the temperature $T$. Using the length change of a ULF during the conformational change $\Delta l = \mathrm{d}L/\mathrm{d}N$ we obtain:

$$\frac{\mathrm{d}L}{\mathrm{d}t} = \Delta l(N_0 - N)k_0 \exp\left(Fx_u/k_BT\right) \tag{2}$$

$$= L_0(\Delta l/l - \varepsilon_{\mathrm{approx}})k_0 \exp\left(Fx_u/k_BT\right), \tag{3}$$

with the length of an ULF $l$, the initial filament length $L_0$ and the strain $\varepsilon_{\mathrm{approx}} = N\Delta l/L_0$ (excluding elastic and entropic contributions). If a filament is stretched at a constant velocity, we can set $\mathrm{d}L/\mathrm{d}t = v$. Solving for the force gives us

$$F = \frac{k_BT}{x_u}\ln\left(\frac{v}{k_0L_0(\Delta l/l - \varepsilon_{\mathrm{approx}})}\right). \tag{4}$$

Elastic contributions play a significant role in the force response of filaments and can be introduced via an elastic correction $\varepsilon_{\mathrm{corr}} = \varepsilon_{\mathrm{approx}} + F/k_{\mathrm{eff}}L_0$ using the effective spring constant of the filament $k_{\mathrm{eff}}$. Comparing this approximate solution to the numerical solution shows good agreement at low to intermediate strains (Supporting Fig. S8).

A simple scaling law can be derived from equation 4 for the force $F_P$ at the onset of the force plateau when the strain is completely caused by elastic deformation, *i.e.* $\varepsilon_{\mathrm{approx}} = 0$. We find

$$F_P = \frac{k_BT}{x_u}\left(\ln(v/L_0) - \ln\left(k_0\Delta l/l\right)\right). \tag{5}$$

This equation can now be used to independently determine $x_u$ and $k_0$ by performing stretching experiments at different velocities and plotting $F_P$ as a function of the normalized velocity $v/L_0$ and estimating $\varepsilon_{\mathrm{max}}$ based on structural data or from the length of the force plateau.

It has been shown previously that the velocity dependence of vimentin stretching could only be reproduced if independent reactions of subsections of the peptide chain were assumed. When we have multiple subunits (for example the three helices that comprise the vimentin monomer) the differential equation 3 becomes a sum of their respective contributions $L_i$ and we get

$$\frac{\mathrm{d}L}{\mathrm{d}t} = \sum_{i}^{3} \frac{\mathrm{d}L_i}{\mathrm{d}t}. \tag{6}$$

This equation cannot be solved without further constrains. However, we can expect that the scaling in equation 5 still holds, since due to the exponential force dependence the element with the largest $x_u$ will dominate at the onset of the plateau. Therefore, equation 5 was fitted to the plateau force as a function of the pulling velocity $v$ normalized by the initial filament length $L_0$.
